# Age-related differences in network structure and dynamic synchrony of cognitive control

**DOI:** 10.1101/2020.10.09.333567

**Authors:** T. Hinault, M. Mijalkov, J.B. Pereira, Giovanni. Volpe, A. Bakker, S.M. Courtney

## Abstract

Cognitive trajectories vary greatly across older individuals, and the neural mechanisms underlying these differences remain poorly understood. Here, we propose a mechanistic framework of cognitive variability in older adults, linking the influence of white matter microstructure on fast and effective communications between brain regions. Using diffusion tensor imaging and electroencephalography, we show that individual differences in white matter network organization are associated with network clustering and efficiency in the alpha and high-gamma bands, and that functional network dynamics partly explain individual cognitive control performance in older adults. We show that older individuals with high versus low structural network clustering differ in task-related network dynamics and cognitive performance. These findings were corroborated by investigating magnetoencephalography networks in an independent dataset. This multimodal brain connectivity framework of individual differences provides a holistic account of how differences in white matter microstructure underlie age-related variability in dynamic network organization and cognitive performance.

## Introduction

As information and events in our daily lives are moving increasingly faster, the ability to control the influence of distractions and switch between different cognitive tasks has never been more important. Aging especially impacts these control processes^1^. However, there are significant inter-individual differences in cognition among older adults, with some individuals showing strong cognitive decline with age, while others maintain similar cognitive performance to that of younger adults^2,3^. Largely understudied so far, the influence of age-related structural brain changes on the temporal dynamics of brain networks, and how this impacts cognitive performance, could provide important insights into the neural basis of individual differences in older adults. This is important as increasing evidence suggests that the best strategy to prevent dementia is to maintain cognitive abilities during aging, which requires identifying the determinants of preserved cognition in older age. In this study, we investigate the interactions between whole-brain microstructural network measures and dynamic functional network organization as a potential source of cognitive variability in healthy aging, in an attempt to provide clues on what network properties are responsible for better cognition in older adults.

Among the different methods used to assess the structural and functional properties of brain networks, graph theory has proven to be a comprehensive approach that is able to characterize the topological organization of areas over the entire brain^4,5^. A graph consists of a series of nodes (e.g., brain regions) connected by edges (e.g., brain connections). Graph analyses can provide useful information regarding the presence of direct connections between brain areas (i.e., the network’s efficiency), or the presence of functionally segregated clusters of brain regions (i.e., the network clustering). Efficiency and clustering are thus central features of brain networks and have been shown to become critically altered during aging in studies using functional magnetic resonance imaging (fMRI), which, although useful, are constrained by the temporal resolution of fMRI^6,7^. Compared to fMRI, magnetoencephalography and electroencephalography (M/EEG) have a higher temporal resolution, allowing assessment of the dynamics of functional networks in greater detail at shorter timescales. However, most studies have averaged network measures over time, instead of assessing how these measures vary as a function of time (but see^8,9^). Averaging network measures could reduce the sensitivity to discover age-related differences, which may potentially depend on critical differences in task-related temporal dynamics^10^. The influence of structural network characteristics on dynamic M/EEG networks might also reveal crucial factors to understand and predict the individual variability of age-related changes in cognitive performance.

Here, we present the results of our investigation regarding the influence of white matter microstructure on the task-related functional communication among multiple brain regions, and whether fast and effective communication can help explain the cognitive variability across older adults. To test this hypothesis, we applied graph theory analyses to both whole-brain EEG synchrony over time and structural integrity measures derived from diffusion tensor imaging (DTI). Our analyses reveal that network dynamics can help explain individual cognitive performance and that both behavioral and task-related functional measures are associated with the preservation of structural network clustering in patterns similar to those observed in young adults. In addition, we replicate our network stability findings and the influence of individual structural network clustering levels in an independent resting-state dataset from the Cambridge Centre for Ageing and Neuroscience dataset (Cam-CAN^11,12^). By combining different neuroimaging modalities with cognitive measures, our study reveals that the underlying structural white matter connections influence dynamic functional connectivity and cognitive performance, providing a mechanistic explanation for the neural basis of the variability in cognitive performance across older adults.

## Results

### Inhibitory deficit in older adults following working memory updating

In this study, 40 young adults and 40 older adults performed an arithmetic verification task while EEG was recorded. In this task (Figure 1A), participants indicated via a gamepad whether the proposed answer for arithmetic equations (e.g., 8 × 4 = 28) was correct given the currently relevant operation (i.e., addition or multiplication) maintained in working memory (WM). Before this verification phase, participants first saw an operation cue i.e. a “+” for addition problems and an “x” for multiplication problems) indicating the relevant arithmetic operation to be maintained in WM. After a pseudo-randomly jittered delay period, a WM cue (“Hold” or “Flip”) was displayed, followed by a second delay period before the presentation of the equation and its proposed solution. The “Hold” cue instructed participants to maintain the cued operation. The “Flip” cue instructed an update to the other, non-cued operation. The operation sign was not displayed with the three numbers during the verification period, to assess the effectiveness of WM maintenance or updating of the cued operation type. Furthermore, “false-related” problems were included to assess inhibitory performance. In these problems the presented, incorrect answer was the correct answer of the other, irrelevant operation type (i.e. correct sum when multiplication was the relevant rule or vice versa; e.g. 8 × 4 = 12). Arithmetic interference refers to longer reaction time for false-related relative to false-unrelated problems (i.e., the proposed solution is false regardless of the currently relevant rule). We summarize the results of the cognitive task briefly here (see details in^13^). In young adults, arithmetic interference was reduced following operation updating (“Flip trials) relative to when the arithmetic operation was actively maintained throughout the trial (“Hold” trials; Table 1). Older adults differed from young adults in that they showed no reduction of arithmetic interference following operation updating (Age × Cue × Problem interaction, p < .001). These results were interpreted as reflecting a facilitation of inhibition of irrelevant arithmetic facts following WM updating in young adults that is less effective in older adults. Because the main difference between age groups was the effect of the WM cue (Flip vs. Hold) on behavioral performance, we specifically investigated the time period immediately following the WM cue onset in EEG data.

**Table 1.** Participants’ characteristics by age group.

| Variables | Young adults | Older adults | *F* | *P* |
| --- | --- | --- | --- | --- |
| *N* | 40 | 40 | - | - |
| Females:males | 27:13 | 27:13 | - | - |
| Age in years | 23 (4.4) | 71 (5.8) | - | - |
| Years of Education | 16 (1.4) | 16 (2.0) | 0.02 | 0.899 |
| Montréal Cognitive Assessment (MOCA) | - | 28.6 (1.2) | - | - |
| Behavioral interference (overall) | 19.88 (87.4) | 43.03 (88.4) | 1.398 | 0.241 |
| Behavioral interference (following maintenance) | 76.65 (111.3) | 37.83 (127.9) | 2.140 | 0.148 |
| **Behavioral interference (following updating)** | **-36.94 (107.0)** | **48.22 (124.2)** | **10.792** | **0.002** |
| **DTI: clustering coefficient** | **0.042 (0.01)** | **0.030 (0.01)** | **45.653** | **<0.001** |
| DTI: average degree | 54.72 (0.8) | 54.45 (0.9) | 1.928 | 0.169 |
| **DTI: efficiency** | **0.31 (0.04)** | **0.28 (0.02)** | **23.326** | **<0.001** |
| **EEG Alpha clustering variability** | **0.01 (0.01)** | **0.02 (0.02)** | **4.833** | **0.031** |
| **EEG High-gamma efficiency variability** | **0.01 (0.01)** | **0.02 (0.01)** | **6.045** | **0.016** |
| **Grey matter volume (mm<sup>3</sup>)** | **618355 (61140)** | **516195 (36503)** | **8.233** | **<0.001** |

*Notes: Average and standard deviations (SD). Behavioral interference refers the difference in reaction time between false-related relative trials and false-unrelated trials in milliseconds. See Hinault et al., 2020, for additional information.*

**Figure 1.**
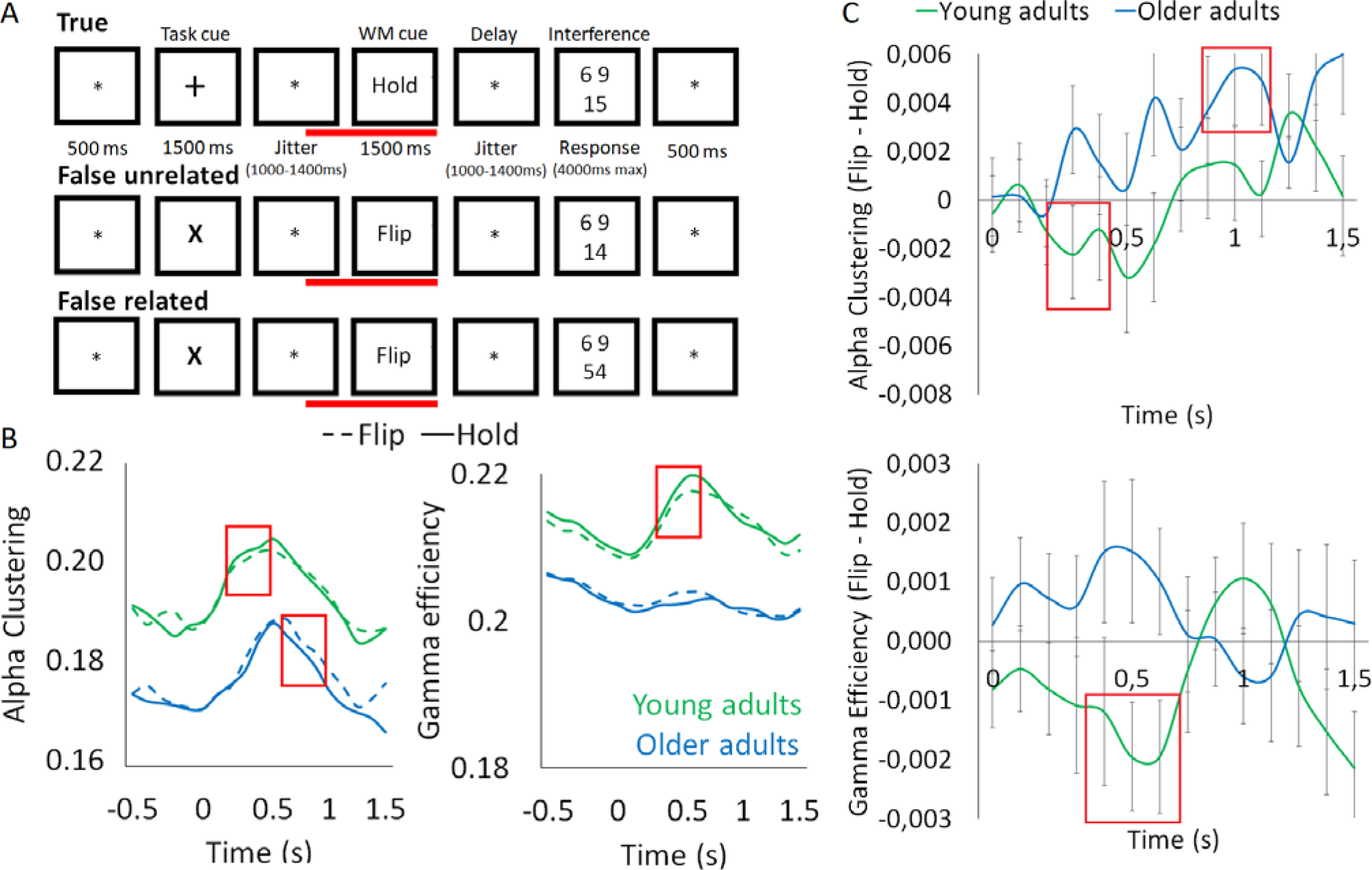
[A] The experimental paradigm was identical to our previous studies (see Hinault et al., 2019, for detailed information regarding the task parameters and the list of stimuli). The red bar indicates the beginning (−500ms before the WM cue onset) and end (1500ms after the WM cue onset) of the sliding windows for the dynamic connectivity analysis. [B] Alpha clustering and high-gamma efficiency metrics for young and older adults following either the maintenance (Hold) or the updating (Flip) cue. Significant differences between age groups are highlighted with red boxes. [C] Difference in alpha clustering and high-gamma efficiency metrics (with standard error) for young and older adults following between the updating (Flip) and the maintenance (Hold) cues. Significant between-condition differences in each group are highlighted with red boxes.

### Age-related changes in dynamic oscillatory networks

In line with previous findings,^13,14^ differences in functional network metrics between young and older adults were investigated in the alpha and high-gamma bands before and after the onset of the WM cue (Figure 1B, 1C; see videos in supplementary information). To test the hypothesis that network dynamics can help explain individual cognitive performance in older adults, we calculated brain networks’ clustering and efficiency. Clustering reflects the prevalence of clusters of brain regions that are strongly connected to each other (but weakly connected to regions in other clusters). Efficiency reflects the direct connections between brain regions (i.e., networks with more direct connections are considered to be more efficient). The graph measures in this study were calculated based on the phase-locking value (PLV) matrix of 200ms sliding time windows before and after the display of the WM cue (19 time windows from −500ms to 1500ms after the onset of the WM cue) for each cortical region of the Desikan atlas^15^, paired with every other region. Interactions among factors of interest (Age, WM cue, Time window) were observed in both alpha and high-gamma bands (Figure 2A). In the alpha band, this interaction involved the network clustering coefficient (*F*(18,1404)=1.924, *p*=.03). Clustering differences were observed between Flip and Hold trials from 200 to 400ms following cue onset in young adults (less clustering for Flip than Hold) and the opposite Cue effect (greater clustering for Flip than Hold) in the 900 to 1100ms time window in older adults. In the high-gamma band, the interaction was observed in network efficiency rather than clustering (*F*(18,1404)=1.794, *p*=.021). Network efficiency was lower for Flip than Hold trials in young adults, between 500 and 700ms following cue onset, but there was no cue-related difference in efficiency in older adults. No main effects were observed. All results were corrected for multiple comparisons using the False Discovery Rate (FDR).

**Figure 2.**
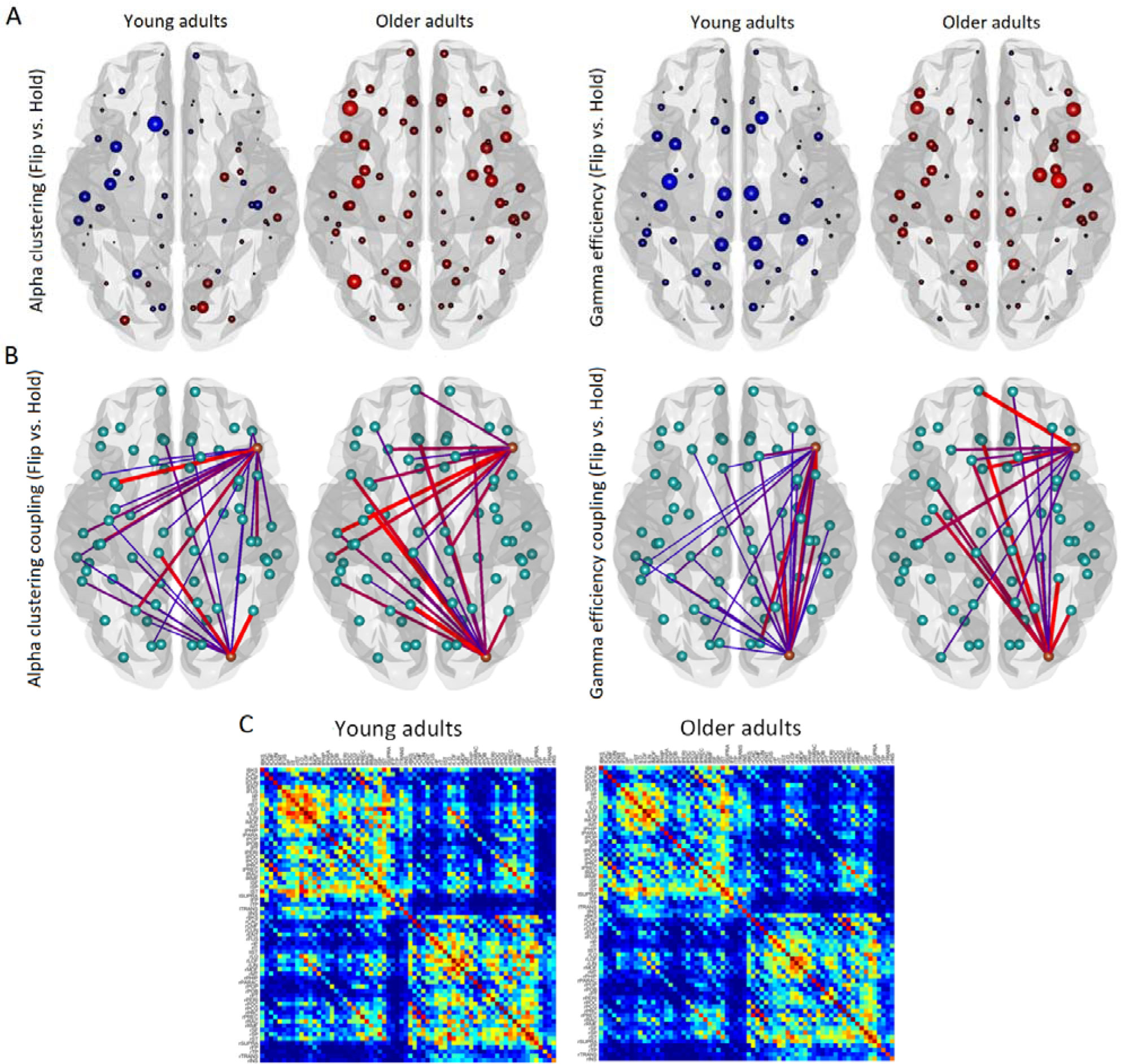
[A] Difference in alpha clustering coefficient and high-gamma efficiency, at each node, between Flip and Hold cues for young and older adults. Size and color indicate nodal values, showing the update-related reduction (blue, Flip < Hold) of alpha clustering and high-gamma efficiency, primarily in young adults, and the update-related increase (red, Flip > Hold) of alpha clustering and high-gamma efficiency, primarily in older adults. [B] Differences in alpha and high-gamma network couplings originating from the right IFG and occipital lobe for Flip vs. Hold cues in young and older adults. Flip > Hold shown in red and Flip < Hold shown in blue. [C] Structural connectivity matrices in young and older adults, average FA value of tracts connecting each ROIs of regions for the Desikan atlas.

Network clustering in the alpha band and network efficiency in the high-gamma band both showed significantly larger variability across time windows, not time-locked to any task event, in older adults than in young adults (*p*=0.031, *p*=0.016, respectively). This larger variability indicates lower network stability (see also^16^). In addition to these global network metrics, consistent results were observed at the nodal (region of interest) level: alpha clustering and high-gamma efficiency effects were observed specifically in edges involving the right IFG (Figure 2B), which has previously been implicated in inhibitory control^17–19^. No effects involving other metrics or brain regions were observed following FDR corrections.

Within the time windows that showed a significant difference between Flip and Hold in either the older or younger groups, we examined correlations between individual differences in the magnitude of that effect and inhibitory control. Among the older adults, the average magnitude of the difference was correlated with behavioral interference for both alpha clustering and high-gamma efficiency in the time windows showing a difference in young adults (*r*=0.407, *p*=0.009; and *r*=0.413, *p*=0.008, respectively). The results (Figure 3A), therefore, indicate that greater Flip – Hold modulations (the opposite of young adults) of alpha clustering and high-gamma efficiency are associated with larger interference effects (i.e. worse cognitive control). In other words, young-like update-related modulations of functional network organization appears to be necessary for optimal task performance. In order to better understand the relationship between functional network metrics and cognitive performance, least-square linear regressions were performed separately for young and older adults, and separately for alpha clustering and high-gamma efficiency metrics, testing for correlations with behavioral interference. Results revealed that high-gamma efficiency (in the time window showing significant differences between Flip and Hold WM cue within that group) explained a significant amount of the behavioral interference variance in both young (R-2=.359, *p*=.027) and older adults (R-2=.407, *p*=.009). In older adults but not young adults, alpha clustering was also found to explain a significant amount of the variance in behavioral interference (R-2=.413, *p*=.008).

**Figure 3.**
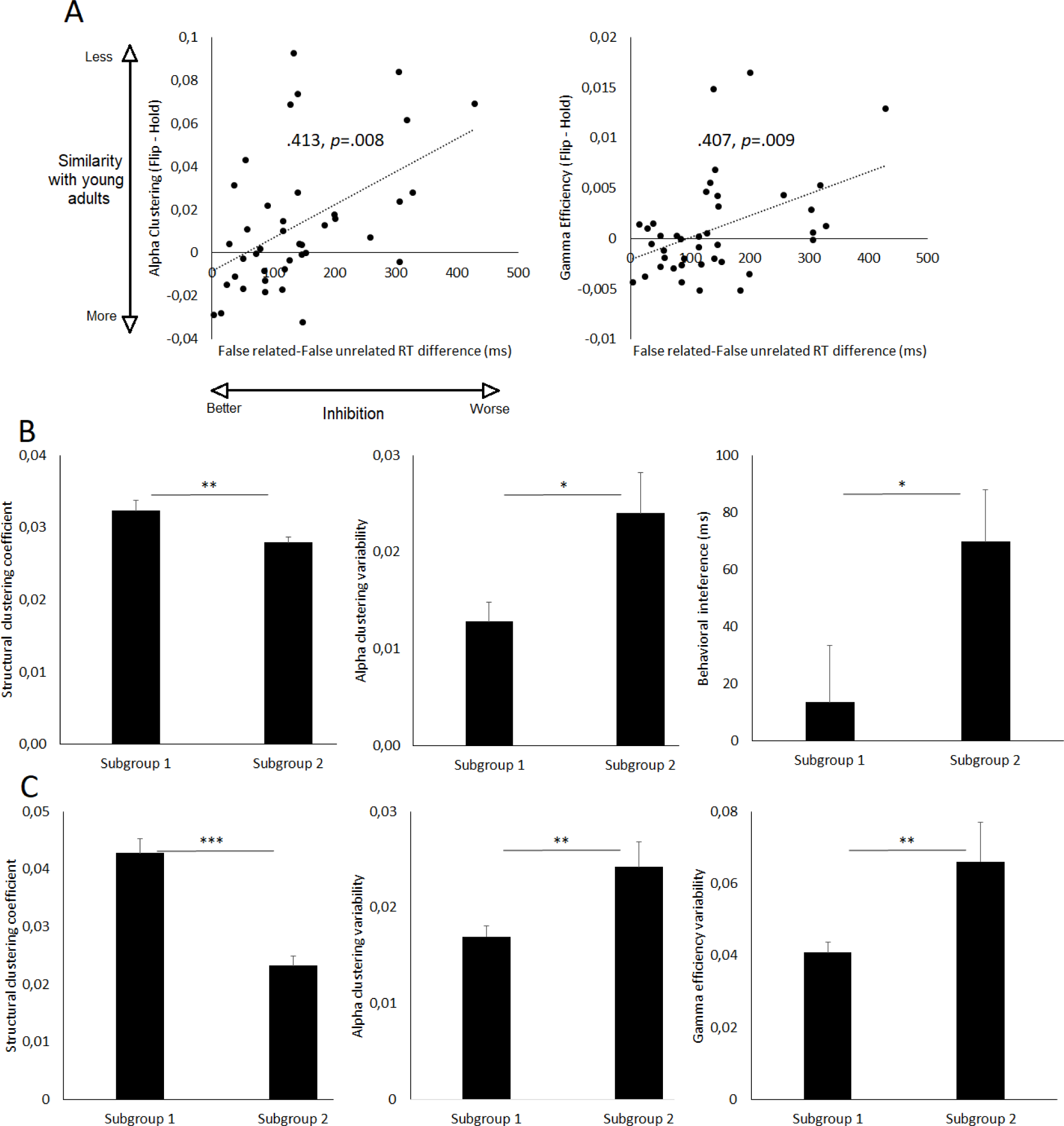
[A] Correlation of alpha clustering and high-gamma efficiency task-related effects with behavioral performance in older adults. Increased Flip – Hold differences (opposite relative to young adults) in the time windows showing significant update-related modulations are associated with larger behavioral interference. [B] DTI clustering subgroup differences in the EEG dataset following data-driven subgroup definition based on structural network clustering. [C] Subgroup differences in the MEG dataset following data-driven subgroup definition based on structural network clustering. Significant differences are highlighted, *p<0.5, \*\**p*<0.1, \*\*\**p*<0.001

### Structural network metrics decrease in older adults

Individual structural connectivity matrices from DTI data were defined based on the average fractional anisotropy (FA) values of tracts connecting cortical regions of the Desikan atlas to investigate differences in structural network integrity (Figure 2C). Comparison of DTI network metrics reveal that young and older participants differ in clustering coefficient levels and efficiency (*p*s<0.001), with both lower clustering and lower efficiency in older adults relative to young adults. No effect of age on any other metric was observed (*F*s<1.5). In contrast with functional network metrics, regression analyses did not reveal a significant amount of variance in behavioral performance that could be explained by a linear relationship with structural network metrics. However, in older adults but not in young adults, structural clustering was positively correlated with larger alpha clustering difference between Flip and Hold (.441, *p*<0.004). In other words, preserved structural network clustering was associated with more young-like update-related modulations of alpha network clustering.

### Data-driven subgroup definition

Given the larger variance in both cognitive and brain measures among older adults compared to young adults^20,21^ and our previous findings on the association between functional and structural connectivity^8,13^, we considered whether there might exist distinct subgroups of older adults driven by white matter microstructural network organization. Such a finding could be important if it indicates a potential biomarker of distinct dynamic network activity and cognitive trajectories with advancing age. To this end, data-driven subgroup definition analyses were conducted based on structural network clustering and efficiency. No viable subgroups for statistical analyses were observed in young adults (32 and 8 individuals in each subgroup for clustering, 38 and 2 individuals in each subgroup for efficiency), which could partially reflect the lower within-group functional variability relative to older adults (*p*s<0.03).

In the older adults group, using the structural clustering metric, two subgroups were identified (19 individuals with higher clustering and 21 individuals with lower clustering; see Table 2, Figure 3B). While the subgroups were similar in age, grey matter volume, cardiovascular fitness, level of education, and high-gamma efficiency, significant differences were observed in behavioral interference and alpha clustering. The subgroup with lower structural clustering showed larger behavioral interference and alpha clustering coefficient in the time window showing significant task-related modulation, compared to the subgroup with higher structural clustering. The variability of alpha network clustering over time is also larger in the subgroup showing lower structural clustering coefficient. Relative to young adults, the older adults’ subgroup with low structural clustering shows larger behavioral interference following updating (*p*<0.001) while the subgroup with higher structural clustering shows no difference (*p*>0.24), suggesting preserved facilitation of inhibition abilities. These subgroup results suggest that older individuals in one subgroup show more young-like structural network clustering, task-related modulations of functional network clustering, stability of functional clustering, and behavioral interference control, relative to the other subgroup that shows greater differences from young adults.

**Table 2.** Differences between subgroups of older adults with different structural network metrics based on hierarchical clustering analyses (standard deviation). The McAuley equation was used to assess cardiorespiratory fitness (McAuley et al., 2012). The equation is based on age, sex, body mass index, resting heart rate, and self-reported physical activity level.

| Variables | Subgroup1 | Subgroup2 | F | p |
| --- | --- | --- | --- | --- |
| *Clustering coefficient* |  |  |  |  |
| N | 19 | 21 | - | - |
| Females:males | 13:6 | 14:7 | 1.567 | 0.218 |
| Age in years | 71 (5.7) | 70 (5.9) | 0.045 | 0.832 |
| Years of education | 15 (1.9) | 16 (2.0) | 3.288 | 0.078 |
| Montréal Cognitive Assessment (MOCA) | 28.58 (1.2) | 28.67 (1.2) | 0.055 | 0.816 |
| **Behavioral interference (overall)** | **13.52 (86.3)** | **69.72 (83.5)** | **4.375** | **0.043** |
| Behavioral interference (following maintenance) | 28.91 (143.3) | 45.91 (115.2) | 0.172 | 0.680 |
| **Behavioral interference (following updating)** | **-1.87 (98.3)** | **93.53 (129.8)** | **6.754** | **0.013** |
| **DTI: clustering coefficient** | **0.032 (0.01)** | **0.027 (0.01)** | **7.959** | **0.008** |
| DTI: average degree | 54.45 (0.9) | 54.46 (0.8) | 0.001 | 0.974 |
| DTI: efficiency | 0.29 (0.1) | 0.28 (0.1) | 3.650 | 0.064 |
| **EEG Alpha clustering variability** | **0.01 (0.1)** | **0.02 (0.1)** | **5.316** | **0.027** |
| **EEG Alpha clustering coefficient (Flip-Hold, 500-700ms)** | **-0.01 (0.1)** | **0.01 (0.1)** | **7.470** | **0.009** |
| EEG High-gamma efficiency variability | 0.01 (0.1) | 0.01 (0.1) | 0.684 | 0.518 |
| Grey matter volume (mm<sup>3</sup>) | 527338<br>(35269) | 506114<br>(35414) | 3.597 | 0.066 |
| Body mass index | 26.62 (5.5) | 27.13 (5.5) | 0.085 | 0.772 |
| Macaley score | 9.04 (1.7) | 8.26 (1.7) | 2.073 | 0.158 |
| *Efficiency* |  |  |  |  |
| N | 17 | 23 | - | - |
| Females:males | 11:6 | 16:7 | 0.876 | 0.355 |
| **Age in years** | **68 (4.3)** | **73 (5.8)** | **9.288** | **0.004** |
| Year of education | 15 (1.8) | 15 (2.2) | 0.001 | 0.976 |
| Montréal Cognitive Assessment (MOCA) | 28.53 (1.1) | 28.70 (1.2) | 0.193 | 0.663 |
| Behavioral interference (overall) | 32.45 (105.6) | 50.84 (74.9) | 0.416 | 0.523 |
| Behavioral interference (following maintenance) | 24.35 (126.7) | 47.80 (130.7) | 0.323 | 0.573 |
| Behavioral interference (following updating) | 40.54 (128.2) | 53.88 (123.7) | 0.110 | 0.742 |
| **DTI: clustering coefficient** | **0.032 (0.1)** | **0.028 (0.1)** | **7.852** | **0.008** |
| DTI: average degree | 54.57 (1.1) | 54.36 (0.7) | 0.601 | 0.443 |
| **DTI: efficiency** | **0.29 (0.2)** | **0.27 (0.1)** | **25.388** | **<0.001** |
| EEG Alpha clustering variability | 0.02 (0.01) | 0.02 (0.2) | 1.415 | 0.242 |
| EEG Alpha clustering coefficient (Flip-Hold, 500-700ms) | 0.01 (0.03) | 0.01 (0.02) | 0.017 | 0.898 |
| EEG High-Gamma efficiency variability | 0.01 (0.1) | 0.01 (0.1) | 0.111 | 0.833 |
| **Grey matter volume (mm<sup>3</sup>)** | **532702**<br>**(34904)** | **503994**<br>**(33315)** | **6.971** | **0.012** |
| Body mass index | 26.00 (4.9) | 27.54 (5.8) | 0.670 | 0.383 |
| Macaley score | 9.11 (1.7) | 8.27 (1.7) | 0.099 | 0.134 |

When analyses were applied to assess structural efficiency in the older adults, two subgroups of similar size were identified (23 and 17 individuals in each subgroup, see Table 2). However, aside from the structural network measures, these subgroups differed only in age and grey matter volume. Participants in the lower structural efficiency subgroup were older and showed greater grey matter atrophy. No significant differences in behavioral performance or functional network metrics were observed. Adopting a reversed approach, we then assessed whether data-driven subgroup definition analyses applied to EEG network metrics would reveal differences in behavioral performance or structural network metrics. No viable cluster was observed with alpha clustering (34 and 6 individuals in each subgroup) or high-gamma efficiency (37 and 3 individuals in each subgroup) metrics, consistent with the interpretation of a specific influence of structural clustering on dynamic networks and cognitive performance.

In summary, the investigation of multimodal network characteristics in putatively healthy older adults revealed changes in network clustering and efficiency across modalities, which were related to cognitive performance. In addition to altered clustering and efficiency of the structural and task-related functional networks, a larger variability across time windows was also observed in older adults relative to young adults, indicating lower functional network stability. A data-driven subgroup definition analysis based on individual structural network clustering levels in the older adults revealed that it could account for individual differences in functional network clustering properties and behavioral performance. Importantly, the same procedure applied to individual structural efficiency revealed only differences of age and grey matter atrophy, and no clearly distinct subgroups were observed when applying the same procedure to EEG network metrics, suggesting a critical role for the preservation of structural clustering in the maintenance of young-like functional network activity and cognitive performance.

### Confirmation from an independent dataset

In order to determine the generalizability of our findings, we analyzed the behavioral, DTI and resting state MEG data from the publicly available Cam-CAN dataset^11,12^. Young and older participants were randomly selected from the Cam-CAN database to match our EEG groups in age and gender balance (see Table 3). Analysis of the DTI data showed that young and older participants differed in efficiency (*p*<0.001) and clustering coefficient (*p*<0.003) metrics. When analyzing resting state MEG data, significant differences between age groups were observed in the variability across time of alpha clustering and high-gamma network efficiency (Table 3, Figure 3C). No main effect of age, or interaction involving any other frequency or metric was observed. Also consistent with findings observed in the EEG dataset, regression analyses revealed that the variability of alpha clustering across time explained a significant amount of the working memory performance variance in older adults (R-2=.139, *p*=.01), but not in young adults. This result was not observed with high-gamma efficiency or with structural network metrics, or with other cognitive measures.

**Table 3.** Characteristics of the participants from the Cam-CAN dataset, and subgroups of older adults defined through hierarchical clustering analyses based on individual levels of structural network clustering coefficient (standard deviation).

| Variables | Young adults | Older adults | *F* | *p* |
| --- | --- | --- | --- | --- |
| *N* | 47 | 47 | - | - |
| Females:males | 30:17 | 30:17 | - | - |
| Age in years | 26 (2.0) | 73 (2.7) | - | - |
| **Years of Education** | **16 (2.8)** | **13 (4.5)** | **12.455** | **0.001** |
| **Mini Mental State Evaluation (MMSE)** | **29.51 (0.9)** | **28.43 (1.2)** | **25.653** | **<0.001** |
| **Visual short term memory (accuracy)** | **0.49 (0.1)** | **0.42 (0.1)** | **21.192** | **<0.001** |
| **Hotel task (seconds)** | **223.24 (122.1)** | **346.43 (174.8)** | **15.232** | **<0.001** |
| **Cattell score** | **37.81 (3.6)** | **26.09 (5.5)** | **150.152** | **<0.001** |
| **DTI: clustering coefficient** | **0.06 (0.1)** | **0.04 (0.1)** | **9.154** | **0.003** |
| **DTI: average degree** | **55.60 (0.9)** | **53.76 (5.3)** | **5.485** | **0.021** |
| **DTI: efficiency** | **0.35 (0.1)** | **0.31 (0.1)** | **30.712** | **<0.001** |
| **MEG alpha clustering variability** | **0.14 (0.1)** | **0.20 (0.1)** | **14.231** | **<0.001** |
| **MEG High-gamma efficiency variability** | **0.003 (0.01)** | **0.006 (0.01)** | **26.257** | **<0.001** |
| **Grey matter volume (mm<sup>3</sup>)** | **700319 (64069)** | **588960 (49434)** | **89.002** | **<0.001** |
| *Older adults' subgroups (DTI clustering coefficient)* |  |  |  |  |
| Variables | Subgroup1 | Subgroup2 | *F* | *p* |
| *N* | 30 | 17 | - | - |
| Females:males | 21:9 | 9:8 | 1.349 | 0.252 |
| Age in years | 73 (2.4) | 73 (3.0) | 0.986 | 0.326 |
| Years of Education | 13 (4.9) | 13 (3.9) | 0.314 | 0.578 |
| Mini Mental State Evaluation (MMSE) | 28.37 (0.2) | 28.53 (0.3) | 0.198 | 0.658 |
| Visual short term memory | 0.43 (0.1) | 0.41 (0.1) | 1.489 | 0.229 |
| Hotel task (seconds) | 324.24 (31.7) | 385.59 (42.5) | 1.347 | 0.252 |
| Cattell score | 25.97 (1.1) | 26.29 (1.0) | 0.038 | 0.846 |
| **DTI: clustering coefficient** | **0.043 (0.01)** | **0.023(0.01)** | **28.511** | **<0.001** |
| **DTI: average degree** | **54.92 (1.1)** | **51.71 (8.5)** | **4.273** | **0.044** |
| **DTI: efficiency** | **0.328 (0.01)** | **0.276 (0.07)** | **15.515** | **<0.001** |
| **MEG alpha clustering variability** | **0.017 (0.01)** | **0.024 (0.01)** | **8.668** | **0.005** |
| **MEG High-gamma efficiency variability** | **0.041 (0.01)** | **0.066 (0.01)** | **7.665** | **0.008** |
| Grey matter volume (mm<sup>3</sup>) | 594751 (44057) | 578741 (57738) | 1.142 | 0.291 |

Similarly to the EEG dataset, data-driven subgroup identification based on structural clustering revealed two subgroups in older adults (17 and 30 participants; see subgroups’ characteristics in Table 3). While subgroups showed similar age, grey matter volume, MMSE score and level of education, significant differences were observed in the variability of both alpha clustering (as in our EEG subgroup data) and high-gamma efficiency. No viable subgroup for statistical analyses was observed in young adults (43 and 4 individuals in the subgroups) or when using MEG metrics (41 and 6 individuals in the subgroups). This replicates our findings with our own data set that older individuals with lower structural network clustering coefficient show less stable functional network clustering. In addition, the Cam-CAN dataset also showed that lower structural clustering was associated with less stable functional network efficiency.

## Discussion

The present study demonstrates the critical importance of considering dynamic M/EEG network characteristics and their associations with other imaging modalities to understand cognitive variability in older adults. We investigated whether task-related dynamic network activity might be critically influenced by individual structural network properties, and might partly explain cognitive performance in older adults with no known clinically significant neuropathology. To this end, we adopted a multimodal approach (EEG, MEG, and DTI data; see supplementary information for fMRI and biological markers’ results) and we applied graph theory analyses to quantify individual structural network clustering levels and their relationship to dynamic functional networks observed during the completion of a task specifically designed to engage cognitive control processes. Results revealed that in older adults, lower structural clustering is associated with changes in clustering and efficiency of functional network task-related dynamics, and lower stability of network metrics across time windows, relative to individuals with structural network characteristics more similar to younger individuals. These differences were also associated with individual differences in cognitive performance. Furthermore, cross-validation of our results in an independent sample with the Cam-CAN dataset provides important confirmation regarding the reliability of these conclusions.

### Dynamic networks and cognitive performance

We focused on the systems underlying cognitive control, and thus investigated dynamic connectivity following WM updating cue onset. Previous studies found that healthy older adults do not generally show facilitation of inhibition following WM updating, in contrast to young adults^8,13^. Recent work highlights the crucial role of alpha activity in the prioritization or de-prioritization of working memory representations^22,23^. This process of WM de-prioritization appears to prime subsequent inhibitory control in young adults. Results in the current study revealed that following WM updating cue onset, relative to the maintenance cue, both alpha clustering and high-gamma efficiency are decreased in young adults. This result suggests an interruption of the previously established functional clustering and the engagement of new, distant pathways and brain regions across clusters during updating. In older adults, these modulations of alpha clustering are delayed and reversed, resulting in increased clustering following the updating cue relative to the maintenance cue, and no modulation of high-gamma efficiency. The inability to quickly and efficiently interrupt the currently engaged sub-network to engage new network regions associated with the other arithmetic operation likely results in lower WM updating ability, and thus was predictive of individual differences in behavioral interference. Nodal analyses also furthered our understanding of the main regions engaged during task performance. Our previous work^13^ reported significant differences between young and older adults in IFG-occipital lobe coupling. The current analyses support and extend those previous results by revealing that edges involving the right IFG are the part of the whole-brain network that is most influenced following the WM cue. These results support the idea that areas in the right IFG play a role not only in inhibiting motor action, but also in interpreting stimuli in the context of current task rules and in rapidly directing multiple changes in functional connectivity^24^.

### Age-related DTI network changes and effects on dynamic functional networks

Age-related modulations of network clustering and efficiency were also observed when investigating DTI structural networks. This suggests a strong relationship between the whole-brain dynamics and the underlying white matter network across the lifespan. In the opposite direction than what has been reported from childhood to adulthood^25^, we observe an overall spatiotemporal connectivity reorganization with aging consisting of reduced clustering (i.e., differentiation into highly connected subnetworks relative to the rest of the network) and efficiency (i.e., highly connected network “hubs” able to reach distant regions^6,26^) of task-engaged networks at both functional and structural levels. Findings also replicate the previously reported association between functional variability and reduced structural integrity in older adults^27^. This suggests that reduced structural connectivity impairs the stability of communications between brain regions.

### Structural network properties underlie functional network organization and cognitive performance

With aging, an overall reduction of integrity of white matter tracts has previously been reported^28^, with long white matter connections being more vulnerable to age-related alterations^29^. These age-related changes lead to decreased structural network efficiency and clustering into subnetworks. The variation of such evolution of structural network clustering and efficiency between individuals appears to be a major determinant of dynamic network activity and cognitive performance with aging. Data-driven subgroup definition analyses reveal that structural network clustering is critically associated with EEG network activity and cognitive performance. This pattern is not observed when defining subgroups based on structural efficiency or functional network measures, suggesting that the preservation of structural network clustering specifically is crucial for the maintenance of functional connectivity dynamics and the ability to suppress interference as effectively as young adults. Results are consistent with our previous studies in showing how structural-functional couplings can account for individual differences in cognitive performance during healthy aging^8,13^, and they further show how whole-brain dynamics can provide data-driven elements to distinguish different cognitive performance patterns within older adults. Interestingly, while only structural network metrics led to the identification of subgroups in older adults that differ in functional network metrics and behavioral performance, only functional network metrics were found to explain a significant amount of the variance associated with behavioral interference resolution. This pattern of result suggests that individual differences in structural network metrics can be used to identify different subgroups varying on both functional network metrics and behavioral performance, while functional network metrics appear to be the more proximal cause of impairments in cognitive performance.

### Corroborating findings in an independent dataset

Additional analyses were performed using the Cam-CAN dataset to determine the degree of generalization of results and the subgroup identification method. Similar to our EEG dataset, differences in structural network clustering were observed between young and older participants, and were then used to identify two subgroups within the older adults that differed in MEG resting state dynamic networks. Consistent with our EEG dataset, older adults in the Cam-CAN dataset with lower structural network clustering showed lower high-gamma network efficiency and larger variability of both alpha clustering and high-gamma clustering relative to the other subgroup. Interestingly, unlike in our EEG dataset, no difference in cognitive performance was observed between the subgroups. This could reflect the lower sensitivity or specificity of the cognitive tests used, as none of the tests in the Cam-CAN dataset specifically targets working memory updating or inhibitory processes. The results confirm the findings from our EEG dataset in highlighting the influence of structural network properties on dynamic network activities.

### Theoretical considerations

Our results differ from the disconnection hypothesis of cognitive aging, proposed to account for cognitive alterations following brain alterations that might “disconnect” brain regions^30,31^. Here, subgroups of older adults do not differ in degree or path length of either structural or functional networks, suggesting that the interruption of network activity along specific pathways is not the primary determinant of the cognitive variability and the typical evolution of functional networks with aging. Rather, results show that the decreased structural, clustered organization of white matter architecture leads to changes in functional network efficiency, clustering, and stability over time, resulting in reduced cognitive performance. This result is in line with the tendency toward more random network architecture and activity with age^6,32^. Our findings point to a possible mechanistic account for the individual variability in cognition associated with aging and suggest that the frameworks of cognitive reserve and reduced speed of processing with aging^33–35^ should include dynamic network activity. The present results highlight that fast and effective, task-relevant modulations of communications between brain regions are crucial for cognitive performance. The current research does not directly establish causation, but the results suggest that preserved structural integrity is necessary for preservation of cognitive performance, because changes in structural integrity lead to less stable functional network organization, and changes in the brain areas recruited for cognitive control, and the organization of their functional connections. The results suggest that this change in functional network dynamics then results in diminished cognitive performance. At the clinical level, the results suggest there might be new ways of identifying individuals at risk of cognitive decline based on structural network organization and functional synchrony. Functional couplings could also be targeted by cognitive training programs or oscillatory neural stimulation (e.g., TMS, tACS) protocols to try to specifically influence the dynamic synchrony between brain regions.

## Methods

### Participants

Data is the same as in our previously published report^13^, and includes 40 young adults (18-35 years) and 40 older adults. All participants are right-handed and reported normal or corrected-to-normal vision. Age groups were matched by years of education, gender, and arithmetic fluency. All older adults performed within normal range (i.e., score > 26) on the Montreal Cognitive Assessment (MoCA^36^). No participant reported a history of neurological or cognitive disorders, traumatic brain injury, or major psychiatric disorders. Participants were not on any neurologically/psychiatrically active medication at the time of testing. See^13^ for additional information regarding exclusion criteria and participants’ characteristics.

### MRI acquisition

Individuals were scanned with a 3T Phillips Achieva MRI scanner with a 32-channel head coil. Total scanning time was 25 minutes. Following localizer scans, T1-weighted images were acquired with a 3D MP-RAGE sequence: Field of view: 212 × 212 × 172 mm, sagittal orientation, 1 × 1 × 1 mm voxel size, repetition time (TR): 3000ms, echo time (TE): 2ms, flip angle: 8°. Diffusion whole brain images were acquired with the following parameters: 70 axial slices, slice thickness = 2.2 mm, voxel size = 2.21 × 2.21 × 2.2 mm, TR = 2000ms, TE = 71ms, FOV = 212 × 212 × 154 mm, SENSE factor: 2.5, BW/pixel: 3124.9. Diffusion gradients were applied along 64 noncollinear directions (b = 1000 s/mm^2^). See supplementary information, for acquisition and analyses of resting-state fMRI data.

### Electroencephalography recording and analyses

EEG activity was recorded inside a Faraday cage with 128 electrodes covering the whole scalp (ActiCHamp, Brain Products, Munich, Germany). Artifact and channel rejection (on continuous data), filtering (0.3-100 Hz bandpass, on unepoched data), re-referencing (i.e., using the algebraic average of the left and right mastoid electrodes), time segmentation into epochs, averaging, and source estimation were performed using Brainstorm^37^. In addition, physiological artifacts (e.g., blinks, saccades) were identified and removed through signal-space projection. The CapTrack camera system was used to record the spatial positions of the electrodes on the cap. These positions were co-registered to that of the individual’s anatomy using fiducial points and were used to improve the source reconstruction accuracy. Age-related brain atrophy can introduce changes in the amplitude of event-related activations^38^. Therefore, FreeSurfer^39^ was used to generate cortical surfaces and automatically segment cortical structures from each participant’s T1-weighted anatomical MRI, to account for individual brain atrophy levels during source reconstruction. The EEG forward model was obtained from a symmetric boundary element method (BEM model; OpenMEEG^40^), fitted to the spatial positions of all electrodes. A cortically constrained, sLORETA procedure was applied to estimate the cortical origin of scalp EEG signals, weighted by a sample estimate of sensor noise covariance matrix obtained from the baseline periods (i.e., fixation period before the display of the WM cue), in each of the participants, and used to improve data modelling. This method was selected because of reduced localization error and false positive connectivity relative to other source localization methods^41,42^. The estimated sources were then smoothed (i.e., full width at half maximum: 3mm) and projected into a standard space (i.e., ICBM152 template) for comparisons between groups and individuals, while controlling for differences in native anatomy. This procedure was applied to activity from 500ms before to 1500ms after the onset of the WM cue.

### Time resolved phase-locking value

Phase-locking analyses^43^ were used to determine the functional coupling between regions of interest. PLV estimates the variability of phase differences between two regions across trials. If the phase difference varies little across trials, PLV is close to 1 (i.e., high synchrony between regions) while, with large variability in the phase difference, PLV is close to zero. The range of each frequency band was adjusted based on the individual alpha-peak frequency (IAF) observed at posterior sites (i.e., bilateral parietal, parieto-occipital, and occipital sites). Following previous literature^44^ and based on previous results using this task^13^, the following frequency bands were then considered: Alpha (IAF-2/IAF+2), low-gamma (IAF+15/IAF+30), and high-gamma (IAF+31/IAF+80). To preserve timing information while reducing the dimensionality of the data, PLV was estimated at the subject level, across trials. To further reduce the dimensionality of the data, the first mode of the principal component analysis (PCA) decomposition of the activation time course in each region of interest (ROI) from the Desikan atlas brain parcellation^15^ was used. The first component, rather than mean activity, was selected to reduce signal leakage^45^. The averages of 200ms sliding time windows (50% overlap) were then extracted across the epochs of interest. In order to avoid arbitrary threshold and unconnected nodes, weighted undirected network analyses were used^46^. See our previous publication^13^ for PLV results for separate pairs of regions. The current report instead uses PLV to quantify whole-brain functional network organization. Similar results were obtained when investigating graph measures derived from lagged coherence matrices, suggesting results are not influenced by common source or volume conduction^46^.

### DTI analyses

Preprocessing of diffusion data was done using ExploreDTI^47^ and included the following steps: (a) Images were corrected for eddy current distortion and participant motion; (b) a non-linear least square method was applied for diffusion tensor estimation, and (c) whole brain DTI deterministic tractography was estimated using the following parameters, for each participant: uniform 2 mm resolution, FA threshold of 0.2 (limit: 1), angle threshold of 45°, and fiber length range of 50 – 500mm. The “network analysis” tools in ExploreDTI were used to quantify the FA value of tracts connecting regions of the Desikan atlas, using the individual cortical parcellation from Freesurfer (Figure 2C). The individual connectivity matrix included all groups of traced fibers identifying a putative tract passing through or ending at two ROIs of the atlas, or null values when no fiber was successfully traced.

### Graph theory analyses

Graph analyses were performed with the Braph toolbox^48^. For EEG data, graph measures were calculated based on the average PLV matrix of 200ms sliding time windows (50% overlap) across the epochs of interest and in each frequency band. For each time window, matrices were defined based on the average PLV synchrony between each region of the Desikan atlas and all other regions (i.e., 68 × 68 matrices, see supplementary information). For DTI data, matrices were defined based on the average FA value of tracts connecting each region of the Desikan atlas with all other regions. Weighted network analyses were used.

Clustering and efficiency were selected as the nodal (i.e., specific to each region) and global (i.e., overall network properties) measures to characterize brain network dynamics and structure based on previous work^49^ and reports of reproducibility^50^: The clustering coefficient is the fraction of triangles (i.e., neighbors of a node that are also neighbors of each other) present around a node and measures the degree to which nodes in a graph tend to cluster together in segregated subnetworks; global efficiency is the average of the inverse shortest path length from a node to all other nodes, representing the network integration and processing capacity. In addition, to ensure that reported effects are not global effects affecting every network parameter, we also examined the average degree (i.e., total number of edges connected to each node, averaged across all nodes) and characteristic path length (i.e., average of the path lengths of all nodes). In addition to Age × Time window and Age × WM cue × Time window mixed-design ANOVAs, the standard deviation of the individual variations of graph measures across time windows was calculated to determine the stability of network activity over time, and corrections for multiple comparisons were performed using False Discovery Rate (FDR) corrections. Participants’ age and mean grey matter volume (from the Freesurfer cortical parcellation) were included as covariates in the analyses. Network analyses were conducted at the nodal level for the clustering coefficient and efficiency measures. Whole-brain analyses were conducted with permutation tests (10,000 permutations), comparing young and older adults on the difference between Flip and Hold trials (Flip *minus* Hold) in the alpha and high-gamma bands. Corrections for multiple comparisons were performed using false discovery rate (FDR) corrections^51^.

### Data-driven subgroup definition

Hierarchical clustering analyses were performed to determine whether subgroups of older adults with different behavioral and functional network patterns could be identified. Hierarchical clustering using the between-groups linkage method was applied on the DTI and EEG graph metrics. Variables were standardized into z-scores before cluster creation. The range of number of clusters started at 2 and stopped at 4, only clusters that would divide the group in subgroups with sufficient number of subjects per subgroup (*N*>15) were selected for statistical analyses.

### Cambridge Centre for Ageing and Neuroscience database

The Cam-CAN dataset^11,12^; available at http://www.mrc-cbu.cam.ac.uk/datasets/camcan/) was used to determine whether the time-varying connectivity and structural network results of the EEG dataset could be replicated in an independent dataset. To match the age and gender characteristics of the EEG groups, data from 47 young adults (20-30 years) and 47 older adults (65-75 years), in participants that completed both structural and functional neuroimaging sessions, were selected for analyses. Details on the demographic and behavioral data can be found at: https://camcan-archive.mrc-cbu.cam.ac.uk/dataaccess/. The selected MRI data included T1-weighted, and diffusion images (see supplementary information). Regarding MEG data, approximately 9 minutes of eyes-closed resting-state data were acquired. Digitization of anatomical landmarks (i.e., fiducial points; nasion and left/right preauricular point, as well as additional points on the scalp) was performed for registration of MEG and MRI coordinate systems. Electro-occulogram and electrocardiogram were recorded to capture eye movements and heartbeats, respectively. Preprocessing involved temporal signal space separation (tSSS): 0.98 correlation, 10s window; bad channel correction: ON; motion correction: OFF; 50Hz+harmonics (mains) notch. The same preprocessing steps as EEG data were followed for MEG data. For MEG data, the selected IAF was the average of the alpha peak observed with gradiometers and magnetometers averaged at posterior sites. In addition to blinks, cardiac artifacts were identified and removed through signal-space projection. No noise modeling was used, as no empty-room recording was available. PLV values averaged across 30s sliding time windows during the resting state period were used to define matrices for graph theory analyses.

## Supporting information

AlphaCluster_FlipvsHold_OA_ADview

AlphaCluster_FlipvsHold_YA_ADview

AlphaSubnetworkConnections_FlipvsHold_OA_ADview

AlphaSubnetworkConnections_FlipvsHold_YA_ADview

GammaEfficiency_FlipvsHold_OA_ADview

GammaEfficiency_FlipvsHold_YA_ADview

GammaSubnetworkConnections_FlipvsHold_OA_ADview

GammaSubnetworkConnections_FlipvsHold_YA_ADview

## Acknowledgements

We would like to thank Eda Incekara, Claire Narang, Daniella Needleman, Pranit Singh, and Phillip Sumardi, for their help with data collection and preprocessing. We also thank Carrie Speck for her help with participant recruitment and screening.

This work was supported by the Albstein Research Foundation, the Fahs-Beck fund for Research and Experimentation, and the William and Ella Owens Medical Research Foundation.

Data collection and sharing for this project were partly provided by the Cambridge Centre for Ageing and Neuroscience (Cam-CAN). Cam-CAN funding was provided by the UK Biotechnology and Biological Sciences Research Council (grant number BB/H008217/1), together with support from the UK Medical Research Council and University of Cambridge, UK.

Joana Pereira and Mite Mijalkov are currently supported by grants from the Swedish Research Council (#2018-02201), Hjärnfonden (#FO2019-0289), Alzheimerfonden (#AF-930827) and the Strategic Research Programme in Neuroscience at Karolinska Institutet (Stratneuro Startup Grant).

## Author contributions

Thomas Hinault designed the research, collected, and analyzed the data, and wrote the paper. Joana Pereira, Giovanni Volpe, Arnold Bakker, and Mite Mijalkov helped analyze the data and write the paper. Susan Courtney helped design the research, analyze the data, and write the paper.

## References

1. Diamond, A. Executive Functions. Annual Review of Psychology 64, 135–168 (2013).

2. Lemaire, P. & Hinault, T. Age-related differences in sequential modulations of poorer-strategy effects. Exp Psychol 61, 253–262 (2014).

3. Li, R. et al. Linking Inter-Individual Variability in Functional Brain Connectivity to Cognitive Ability in Elderly Individuals. Front. Aging Neurosci. 9, (2017).

4. Farahani, F. V., Karwowski, W. & Lighthall, N. R. Application of Graph Theory for Identifying Connectivity Patterns in Human Brain Networks: A Systematic Review. Front. Neurosci. 13, 585 (2019).

5. Vecchio, F. et al. “Small World” architecture in brain connectivity and hippocampal volume in Alzheimer’s disease: a study via graph theory from EEG data. Brain Imaging and Behavior 11, 473–485 (2017).

6. Iordan, A. D. et al. Aging and Network Properties: Stability Over Time and Links with Learning during Working Memory Training. Frontiers in Aging Neuroscience 9, (2018).

7. Sala-Llonch, R. et al. Changes in whole-brain functional networks and memory performance in aging. Neurobiology of Aging 35, 2193–2202 (2014).

8. Hinault, T., Larcher, K., Bherer, L., Courtney, S. M. & Dagher, A. Age-related differences in the structural and effective connectivity of cognitive control: a combined fMRI and DTI study of mental arithmetic. Neurobiology of Aging 82, 30–39 (2019).

9. Coquelet, N. et al. The electrophysiological connectome is maintained in healthy elders: a power envelope correlation MEG study. Sci Rep 7, 13984 (2017).

10. Ariza, P. et al. Evaluating the effect of aging on interference resolution with time-varying complex networks analysis. Front. Hum. Neurosci. 9, (2015).

11. Taylor, J. R. et al. The Cambridge Centre for Ageing and Neuroscience (Cam-CAN) data repository: Structural and functional MRI, MEG, and cognitive data from a cross-sectional adult lifespan sample. NeuroImage 144, 262–269 (2017).

12. Shafto, M. A. et al. The Cambridge Centre for Ageing and Neuroscience (Cam-CAN) study protocol: a cross-sectional, lifespan, multidisciplinary examination of healthy cognitive ageing. BMC neurology 14, 204 (2014).

13. Hinault, T., Kraut, M., Bakker, A., Dagher, A. & Courtney, S. M. Disrupted Neural Synchrony Mediates the Relationship between White Matter Integrity and Cognitive Performance in Older Adults. Cereb Cortex 30, 5570–5582 (2020).

14. Hinault, T., Lemaire, P. & Phillips, N. Aging and sequential modulations of poorer strategy effects: An EEG study in arithmetic problem solving. Brain Research 1630, 144–158 (2016).

15. Desikan, R. S. et al. An automated labeling system for subdividing the human cerebral cortex on MRI scans into gyral based regions of interest. NeuroImage 31, 968–980 (2006).

16. Kumral, D. et al. BOLD and EEG signal variability at rest differently relate to aging in the human brain. NeuroImage 207, 116373 (2020).

17. Aron, A. R., Robbins, T. W. & Poldrack, R. A. Inhibition and the right inferior frontal cortex: one decade on. Trends in Cognitive Sciences 18, 177–185 (2014).

18. Hinault, T., Larcher, K., Zazubovits, N., Gotman, J. & Dagher, A. Spatio–temporal patterns of cognitive control revealed with simultaneous electroencephalography and functional magnetic resonance imaging. Human Brain Mapping 40, 80–97 (2019).

19. Montojo, C. A. & Courtney, S. M. Differential Neural Activation for Updating Rule versus Stimulus Information in Working Memory. Neuron 59, 173–182 (2008).

20. Hedden, T. & Gabrieli, J. D. E. Insights into the ageing mind: a view from cognitive neuroscience. Nature Reviews Neuroscience 5, 87–96 (2004).

21. Hultsch, D. F., Strauss, E., Hunter, M. A. & MacDonald, S. W. S. Intraindividual variability, cognition, and aging. in The handbook of aging and cognition, 3rd ed 491–556 (Psychology Press, 2008).

22. Hinault, T., Blacker, K. J., Gormley, M., Anderson, B. A. & Courtney, S. M. Value-driven attentional capture is modulated by the contents of working memory: An EEG study. Cogn Affect Behav Neurosci 19, 253–267 (2019).

23. de Vries, I. E. J., Slagter, H. A. & Olivers, C. N. L. Oscillatory Control over Representational States in Working Memory. Trends in Cognitive Sciences 24, 150–162 (2020).

24. Xu, K. Z. et al. Neural Basis of Cognitive Control over Movement Inhibition: Human fMRI and Primate Electrophysiology Evidence. Neuron 96, 1447–1458.e6 (2017).

25. Vohryzek, J. et al. Dynamic spatiotemporal patterns of brain connectivity reorganize across development. Network Neuroscience 4, 115–133 (2020).

26. López, M. E. et al. MEG Beamformer-Based Reconstructions of Functional Networks in Mild Cognitive Impairment. Front. Aging Neurosci. 9, 107 (2017).

27. Scarapicchia, V., Garcia-Barrera, M., MacDonald, S. & Gawryluk, J. R. Resting State BOLD Variability Is Linked to White Matter Vascular Burden in Healthy Aging but Not in Older Adults With Subjective Cognitive Decline. Front. Hum. Neurosci. 13, 429 (2019).

28. Molloy, C. J., Nugent, S. & Bokde, A. L. W. Alterations in Diffusion Measures of White Matter Integrity Associated with Healthy Aging. J Gerontol A Biol Sci Med Sci doi:10.1093/gerona/glz289.

29. Yang, A. C., Tsai, S.-J., Liu, M.-E., Huang, C.-C. & Lin, C.-P. The Association of Aging with White Matter Integrity and Functional Connectivity Hubs. Front. Aging Neurosci. 8, (2016).

30. Bennett, I. J. & Madden, D. J. Disconnected Aging: Cerebral White Matter Integrity and Age-Related Differences in Cognition. Neuroscience 0, 187–205 (2014).

31. Madden, D. J. et al. Sources of disconnection in neurocognitive aging: cerebral white-matter integrity, resting-state functional connectivity, and white-matter hyperintensity volume. Neurobiology of Aging 54, 199–213 (2017).

32. Miraglia, F., Vecchio, F. & Rossini, P. M. Searching for signs of aging and dementia in EEG through network analysis. Behavioural Brain Research 317, 292–300 (2017).

33. Cabeza, R. et al. Maintenance, reserve and compensation: the cognitive neuroscience of healthy ageing. Nature Reviews Neuroscience 19, 701 (2018).

34. Salthouse, T. A. Influence of processing speed on adult age differences in working memory. Acta Psychologica 79, 155–170 (1992).

35. Stern, Y. et al. Whitepaper: Defining and investigating cognitive reserve, brain reserve, and brain maintenance. Alzheimer’s & Dementia (2018) doi:10.1016/j.jalz.2018.07.219.

36. Nasreddine, Z. S. et al. The Montreal Cognitive Assessment, MoCA: a brief screening tool for mild cognitive impairment. Journal of the American Geriatrics Society 53, 695–699 (2005).

37. Tadel, F., Baillet, S., Mosher, J. C., Pantazis, D. & Leahy, R. M. Brainstorm: A User-Friendly Application for MEG/EEG Analysis. Computational Intelligence and Neuroscience 2011, 1–13 (2011).

38. Moretti, D. V., Paternicò, D., Binetti, G., Zanetti, O. & Frisoni, G. B. EEG upper/low alpha frequency power ratio relates to temporo-parietal brain atrophy and memory performances in mild cognitive impairment. Front. Aging Neurosci. 5, (2013).

39. Fischl, B. FreeSurfer. NeuroImage 62, 774–781 (2012).

40. Gramfort, A., Papadopoulo, T., Olivi, E. & Clerc, M. OpenMEEG: opensource software for quasistatic bioelectromagnetics. BioMedical Engineering OnLine 9, 45 (2010).

41. Kitaura, Y. et al. Functional localization and effective connectivity of cortical theta and alpha oscillatory activity during an attention task. Clin Neurophysiol Pract 2, 193–200 (2017).

42. Pascual-Marqui, R. D. et al. Comparing EEG/MEG neuroimaging methods based on localization error, false positive activity, and false positive connectivity. bioRxiv (2018) doi:10.1101/269753.

43. Lachaux, J.-P., Rodriguez, E., Martinerie, J., Varela, F. J. & others. Measuring phase synchrony in brain signals. Human brain mapping 8, 194–208 (1999).

44. Toppi, J. et al. Different Topological Properties of EEG-Derived Networks Describe Working Memory Phases as Revealed by Graph Theoretical Analysis. Frontiers in Human Neuroscience 11, (2018).

45. Sato, M., Yamashita, O., Sato, M. & Miyawaki, Y. Information spreading by a combination of MEG source estimation and multivariate pattern classification. PLOS ONE 13, e0198806 (2018).

46. Hardmeier, M. et al. Reproducibility of Functional Connectivity and Graph Measures Based on the Phase Lag Index (PLI) and Weighted Phase Lag Index (wPLI) Derived from High Resolution EEG. PLoS ONE 9, e108648 (2014).

47. Leemans, A., Jeurissen, B., Sijbers, J. & Jones, D. K. ExploreDTI: a graphical toolbox for processing, analyzing, and visualizing diffusion MR data. Proc. Intl. Soc. Mag. Reson. Med. 17, 1 (2009).

48. Mijalkov, M. et al. BRAPH: A graph theory software for the analysis of brain connectivity. PLOS ONE 12, e0178798 (2017).

49. Yu, Q. et al. Building an EEG-fMRI Multi-Modal Brain Graph: A Concurrent EEG-fMRI Study. Frontiers in Human Neuroscience 10, (2016).

50. Welton, T., Kent, D. A., Auer, D. P. & Dineen, R. A. Reproducibility of Graph-Theoretic Brain Network Metrics: A Systematic Review. Brain Connectivity 5, 193–202 (2015).

51. Benjamini, Y. & Hochberg, Y. Controlling the False Discovery Rate: A Practical and Powerful Approach to Multiple Testing. Journal of the Royal Statistical Society. Series B (Methodological) 57, 289–300 (1995).

