## Supplementary information for "Age-related differences in network structure and dynamic synchrony of cognitive control"

**Methods:**

**Resting state fMRI data.** FMRI data were acquired while participants performed a 10-minutes resting state session, using a whole brain gradient echo planar imaging sequence sensitive to BOLD contrast in 48 slices (RT: 3,000 ms, ET: 30 ms, 3.5 mm slice thickness, voxel sizes: 3.31 × 3.31 × 3.31 mm, FA: 80°, FOV: 250 × 250 × 159 mm). Data preprocessing was performed using the CONN toolbox^1^ (<http://www.nitrc.org/projects/conn>). The default preprocessing pipeline was applied^2^, including slice-timing correction, realignment of functional images to correct for head motion, high-pass filtering (0.01 Hz) to remove signal drift, normalization into the Montreal Neurological Institute (MNI) stereotaxic space with trilinear interpolation. Images were also convolved spatially with a 3D isotropic Gaussian kernel (8 mm Full Width at Half Maximum [FWHM]) to improve signal-to-noise ratio. Connectivity matrices were defined based on the correlation of the BOLD signal between regions of the Desikan atlas in sliding windows (100s, 50% overlap) across the resting state period.

**Biological data.** CSF data were collected in a subgroup of participants (N:19; see also in^3^). 20 ml CSF was collected in the morning between 8 and 10 am, after an overnight fast, into a 50 ml polypropylene tube. After mixing and centrifugation at 2000 rpm for 15 minutes, 500 ul aliquots of CSF were frozen at -80 °C within 60 minutes of collection. CSF Aβ 42 (picograms/ml) and CSF total tau (picograms/ml) were measured using the Lumipulse G1200 assay (Fujirebio, Malvern, PA). Assays were run in duplicate, and all samples were run in a single batch. Intra-assay coefficient of variation for this assay was 3.1% for Aβ 42 and 4.3% for total tau.

**Cambridge Centre for Ageing and Neuroscience database.** The selected MRI data included T1-weighted (Field of view: 256 × 240 × 192 mm, 1 × 1 × 1 mm voxel size, RT: 2250ms, ET: 900ms, ﬂip angle: 9°), and diffusion (66 axial slices, slice thickness = 2 mm, RT = 9100ms, ET = 104ms, FOV = 192 x 192 x 2 mm) images. Diffusion gradients were applied along 30 noncollinear directions (b = 1000/2000s/mm²). Regarding MEG data, approximately 9 minutes of eyes-closed resting-state data were acquired with a 306-channel Elekta Neuromag Vectorview (102 magnetometers and 204 planar gradiometers), at 1kHz sampling, and 0.03-330 Hz online filter.

**Results:**

**Association with biological markers of pathological aging***.* Measures of Aβ42 (mean: 138) and total Tau levels (mean: 761). There measures are considered as markers of pathological aging, as CSF Aß 42 is used to index amyloid-beta (Aß) protein deposition, while CSF total tau (t-tau) is associated with neurodegeneration^3^. In a subset of our older participants (N=19), we investigated the association between these measures and EEG metrics in time windows that showed significant effects. Results revealed significant correlations between Aβ42 levels and high-gamma efficiency 500 to 700ms following the Flip WM cue (*r*=-.627, *p*=0.004), with lower concentrations associated with larger high-gamma efficiency. Correlations were also observed between Tau levels and alpha clustering coefficient 200 to 400ms following the Flip WM cue onset (*r*=.629, *p*=0.004). Increased Tau levels were associated with higher alpha clustering coefficients (see Figure S1). No significant correlations were observed for the Flip *minus* Hold difference.

*Figure S1. Correlations between EEG gamma efficiency and alpha clustering network characteristics and CSF concentrations of Aβ42 and tau in a selected subgroup of older adults.*

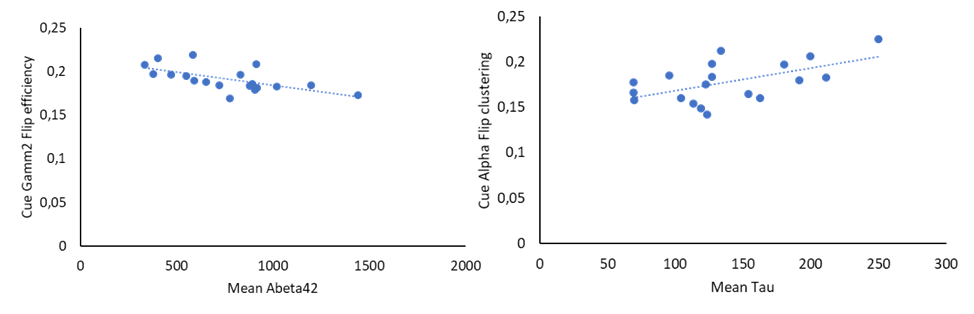

.629, *p*=.004

-.627, *p*=.004

**Electroencephalography analyses.** Following recent work^4^, the relative alpha band power was computed as the ratio between the sum of the alpha power spectrum density (PSD) over the alpha frequency range and the sum of the total PSD over the frequency range in 1-90Hz. Power was computed for each ROI at the source level and alpha frequency range was based on individual posterior alpha peak frequency. Following FDR correction for multiple comparisons, negative correlations were observed in both young and older adults, between alpha power and the variability of clustering network metric across time windows (*p*s<0.009). This result is consistent with the partial association between power and network metrics but does not appear to vary as a function of age.

**Age-related changes of fMRI network metrics**. Network analyses of rs-fMRI data revealed significant differences between young and older adults in clustering coefficient (*p*=.017) and efficiency (*p*=.016), with larger values in older adults relative to young adults. No correlations between rs-fMRI network metrics and behavioral performance were found, however.

**Discussion:**

**Dynamic networks and cognitive performance.** Results on the association between EEG source power and network metrics are in line with previous work^4^ in showing correlations between power spectrum density and network metrics derived from PLV. However, in contrasts with network measures, effects are observed in both young and older adults, suggesting that power analyses might be less sensitive to aging effects than network measures, due to age-related changes in brain structure and function.

**Dynamic networks and biological markers.** Oscillatory networks were also associated with known biological markers of pathological aging. Aβ42 and Tau CSF concentrations were obtained in a subset of the older adults’ group. Results reveal that lower mean Aβ42 concentrations are associated with higher efficiency in the high-gamma EEG network, while clustering of the alpha network is higher with increased Tau concentrations. This pattern is surprising, given that the neurochemical profile of Alzheimer’s disease is defined by higher Tau and lower Aβ42 CSF concentrations^8,9^. Given that only healthy older adults have been recruited, with normal cognitive performance and no known neurological conditions, it is possible that these associations with functional network activity reflect compensatory adjustments at very early stages of neurodegenerative processes. Future studies will investigate how other age-related changes in biological markers of pathological aging contribute to EEG dynamic networks.

**References:**

1.	Whitfield-Gabrieli, S. & Nieto-Castanon, A. Conn: A Functional Connectivity Toolbox for Correlated and Anticorrelated Brain Networks. *Brain Connectivity* **2**, 125–141 (2012).

2.	Nieto-Castanon, A. *Handbook of functional connectivity Magnetic Resonance Imaging methods in CONN*. (Hilbert Press, 2020).

3.	Alm, K. H. *et al.* Medial temporal lobe white matter pathway variability is associated with individual differences in episodic memory in cognitively normal older adults. *Neurobiology of Aging* **87**, 78–88 (2020).

4.	Demuru, M., La Cava, S. M., Pani, S. M. & Fraschini, M. A comparison between power spectral density and network metrics: An EEG study. *Biomedical Signal Processing and Control* **57**, 101760 (2020).

5.	Hinault, T., Larcher, K., Zazubovits, N., Gotman, J. & Dagher, A. Spatio–temporal patterns of cognitive control revealed with simultaneous electroencephalography and functional magnetic resonance imaging. *Human Brain Mapping* **40**, 80–97 (2019).

6.	Labounek, R. *et al.* EEG spatiospectral patterns and their link to fMRI BOLD signal via variable hemodynamic response functions. *Journal of Neuroscience Methods* **318**, 34–46 (2019).

7.	Suárez, L. E., Markello, R. D., Betzel, R. F. & Misic, B. Linking Structure and Function in Macroscale Brain Networks. *Trends in Cognitive Sciences* **24**, 302–315 (2020).

8.	Galasko, D. *et al.* High cerebrospinal fluid tau and low amyloid beta42 levels in the clinical diagnosis of Alzheimer disease and relation to apolipoprotein E genotype. *Arch. Neurol.* **55**, 937–945 (1998).

9.	Sunderland, T. *et al.* Decreased β-Amyloid1-42 and Increased Tau Levels in Cerebrospinal Fluid of Patients With Alzheimer Disease. *JAMA* **289**, 2094–2103 (2003).
